# Obesity-Activated Lung Stomal Cells Promote Myeloid-Lineage Cell Accumulation and Breast Cancer Metastasis

**DOI:** 10.1101/2020.11.23.394387

**Authors:** Lauren E. Hillers-Ziemer, Abbey E. Williams, Amanda Janquart, Caitlin Grogan, Victoria Thompson, Adriana Sanchez, Lisa M. Arendt

**Affiliations:** Program in Cellular and Molecular Biology, University of Wisconsin-Madison, Madison WI 53706, USA; Program in Comparative Biomedical Sciences, University of Wisconsin-Madison, Madison WI 53706, USA; Department of Comparative Biosciences, School of Veterinary Medicine, University of Wisconsin-Madison, Madison WI 53706, USA

**Keywords:** Obesity, breast cancer, metastasis, macrophages, collagen, lung fibroblasts, Met-1, TC2

## Abstract

Obesity is correlated with increased incidence of breast cancer metastasis, however the mechanisms underlying how obesity promotes metastasis are unclear. In a diet-induced obesity mouse model, obesity enhanced lung metastases in both the presence and absence of primary mammary tumors and increased recruitment of myeloid lineage cells into the lungs. In the absence of tumors, obese mice demonstrated increased numbers of myeloid lineage cells and elevated collagen fibers within the lung stroma, reminiscent of pre-metastatic niches formed by primary tumors. Lung stromal cells isolated from obese non-tumor-bearing mice showed increased proliferation, contractility, and expression of extracellular matrix, inflammatory markers, and TGFβ1. Conditioned media from lung stromal cells from obese mice promoted myeloid lineage cell migration *in vitro* in response to CSF2 expression and enhanced invasion of tumor cells. Together, these results suggest that prior to tumor formation, obesity alters the lung microenvironment, creating niches conducive for metastatic growth.

## INTRODUCTION

Global obesity rates, as defined by a body mass index (BMI) greater than 30.0 kg/m^2^, have nearly tripled since 1975; approximately 13% of the world’s population is considered to be obese, including 15% of women (World Health Organization, 2016). Obesity increases the risk for breast cancer in postmenopausal women, as well as premenopausal women who have elevated risk due to heritable factors (Chun et al., 2006; Lahmann et al., 2004; Lauby-Secretan et al., 2016). Regardless of menopausal status, obese breast cancer patients have an enhanced risk for developing distant metastases compared to lean patients (Ewertz et al., 2011; Sestak et al., 2010), particularly to the liver and lungs (Osman and Hennessy, 2015). While the five-year survival rate of metastatic breast cancer patients has significantly increased over the last 30 years (Sundquist et al., 2017), metastasis accounts for the vast majority of breast cancer-related deaths. The mechanisms of how obesity promotes metastatic breast cancer are largely unknown. As obese patients are also at an elevated risk for treatment resistance (Ioannides et al., 2014; Karatas et al., 2017; Sparano et al., 2012), understanding the relationship between obesity and metastasis is vital to develop targeted therapies for obese patients.

Metastasis is a complex process in which tumor cells escape the primary tumor, survive in circulation, extravasate at distal sites, and proliferate in competent organs. Evidence from pre-clinical models has suggested that primary breast tumors promote metastasis through establishment of pre-metastatic niches in potential metastatic organs (Liu and Cao, 2016; Peinado et al., 2017). A major component of pre-metastatic niches are bone marrow-derived myeloid lineage cells, including monocytes, macrophages, neutrophils and myeloid-derived suppressor cells (MDSC). MDSCs are a heterogeneous population of CD11b^+^ myeloid cells classified into 2 subsets: granulocytic MDSCs (gMDSCs), most similar to neutrophils, and monocytic MDSCs (mMDSCs) which resemble monocytes. Expansion of MDSC subtypes is dependent on systemic and microenvironmental cues (Ouzounova et al., 2017; Youn et al., 2008), as MDSCs are absent in healthy individuals (Gabrilovich, 2017) but increase under conditions of obesity (Clements et al., 2018; Okwan-Duodu et al., 2013; Ostrand-Rosenberg and Sinha, 2009). Myeloid lineage cells are thought to aid in the establishment of an environment conducive for metastatic growth through secretion of cytokines, extracellular matrix (ECM) remodeling, and immunosuppression (Liu and Cao, 2016; Swierczak and Pollard, 2020). Although obesity results in recruitment of myeloid lineage cells into obese adipose tissue (Ferrante, 2013), little is known regarding the effects of obesity on the immune populations in distant sites which might contribute to metastasis.

Within the pre-metastatic environment, tumor-secreted factors alter stromal cells, leading to changes in expression of ECM proteins and matrix metalloproteinases (Kong et al., 2019; Liu and Cao, 2016). Studies have suggested that successful metastatic colonization occurs through both structural alterations of the ECM and deposition of new ECM components within pre-metastatic niches (Peinado et al., 2017; Sleeman, 2012). Stromal cells secrete ECM proteins, such as fibronectin, which facilitate tumor cell adhesion and colonization (Paolillo and Schinelli, 2019). Lung stromal cell activation has been observed in other pathological conditions, such as idiopathic pulmonary fibrosis and increased immune cells and serum cytokines have been shown to play a role (Li et al., 2018; Su et al., 2016). Obesity leads to chronic inflammation within adipose tissue, resulting in increased circulating levels of multiple inflammatory cytokines (Dao et al., 2020; Williams et al., 2020). Within the mammary gland, obesity activates adipose-derived stromal cells, promoting mammary tumor progression (Hillers et al., 2018). However, how obesity impacts stromal cells at distant sites has not been examined, and these changes may significantly enhance distal metastatic colonization.

Here, we examined how obesity promotes breast cancer metastasis through activation of the lung microenvironment. We show that obesity enhances metastasis to the lungs both in the presence and absence of primary mammary tumors. Lungs from obese mice demonstrated increased recruitment of myeloid lineage cells prior to and during metastatic growth. In the absence of primary tumors, lung stromal cells isolated from obese mice demonstrated increased proliferation rates, enhanced collagen deposition, and elevated expression of proinflammatory cytokines compared to lung stromal cells from lean mice. Further, conditioned media from lung stromal cells from obese mice enhanced invasion of bone marrow-derived myeloid lineage cells in culture through elevated expression of CSF2. Overall, our findings suggest that obesity activates lung stromal cells prior to tumor formation, leading to increased myeloid lineage cell recruitment, similar to pre-metastatic niche formation by tumor cells during cancer progression. These changes in the lung microenvironment in obesity may contribute to the increased metastatic incidence observed in obese breast cancer patients, as well as other obesity-related cancers.

## RESULTS

### Obesity Increases Mammary Tumor Metastasis

To examine how obesity impacts mammary tumor growth and metastasis, we utilized a high-fat diet (HFD) model of obesity and implanted mammary tumor cell lines into the inguinal mammary glands of mice. Three-week-old female FVB/N mice were fed either a low-fat diet (LFD) or HFD for 16 weeks to induce obesity. HFD-fed mice gained significantly more weight than LFD-fed mice at 7 weeks after starting the HFD (Figure 1A). We have previously shown that after 16 weeks, HFD-fed female FVB/N mice have increased mammary gland weights, larger adipocyte diameters, and elevated numbers of crown-like structures compared with LFD-fed mice (Chamberlin et al., 2017). To investigate how obesity impacts mammary tumor growth, we implanted either Met-1 or TC2 tumor cells into mammary fat pads. Consistent with our previous study (Hillers-Ziemer, 2020), Met-1 and TC2 mammary tumors from HFD-fed mice grew significantly faster than tumors from LFD-fed mice (Figure 1B), indicating that obesity promotes tumor growth.

**Figure 1.**
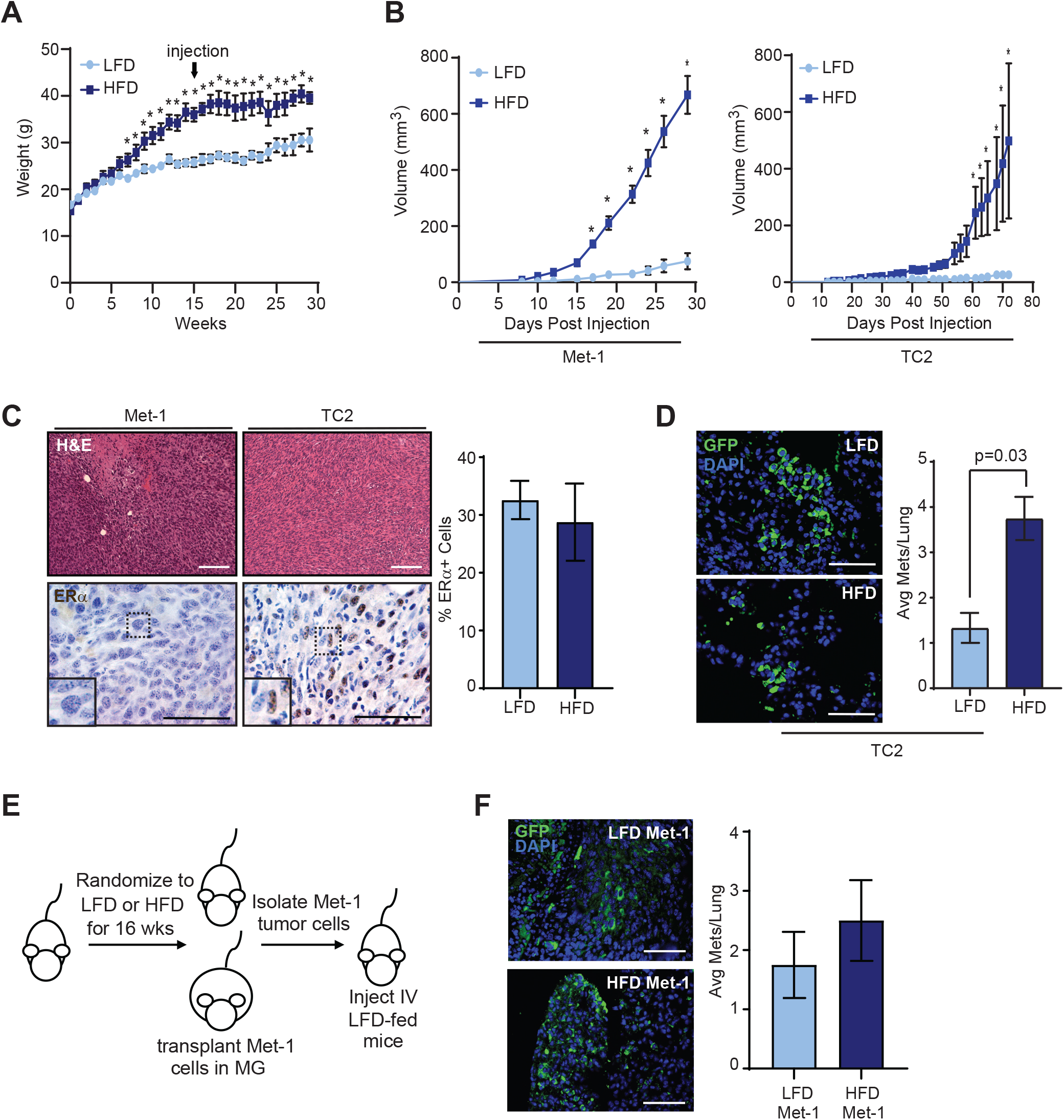
Obesity increases breast cancer metastasis. (A) Weight gain of female mice fed either the low-fat diet (LFD) or high-fat diet (HFD) for 16 weeks prior to tumor cell transplantation (arrow; n=6-8 mice/group). (B) Growth curves of Met-1 or TC2 tumor cells transplanted into mammary glands of LFD or HFD-fed female mice (n=4 mice/group). (C) Representative images of Met-1 and TC2 tumors stained with hematoxylin and eosin (H&E) or immunohistochemistry to detect estrogen receptor alpha (ERα). Quantification of ERα^+^ cells in tumors from LFD or HFD-fed mice (n=4 mice/group). (D) Quantification of metastatic foci of green fluorescent protein (GFP)-expressing TC2 tumor cells in lungs of LFD and HFD-fed mice (n=4 mice/group). (E) Schematic of experiment to inject tumor cells into the tail vein of LFD-fed mice. (F) Quantification of Met-1 metastatic foci in lungs of LFD-fed mice (n=5 mice/group). Magnification bars = 50μm.

Following transplantation, no significant differences were observed histologically among Met-1 or TC2 tumors from LFD and HFD-fed mice. Met-1 tumor cells were derived from a MMTV-PyMT tumor and do not express estrogen receptor alpha (ERα) (Borowsky et al., 2005). Consistent with this previous study, we did not observe ERα expression within tumors of LFD or HFD-fed mice (Figure 1C). In contrast, TC2 tumor cells express ERα in culture and *in vivo* (Barcus et al., 2017) and in tumors from LFD and HFD-fed mice (Figure 1C). No differences were observed in the percentage of ERα-expressing TC2 tumor cells from LFD or HFD-fed mice (Figure 1C). These data indicate that obesity enhances the growth of both ERα^+^ and ERα^−^ tumors.

Clinical evidence suggests that obesity increases the incidence of metastatic breast cancer (Ewertz et al., 2011; Sestak et al., 2010). We have previously shown that HFD-fed mice orthotopically transplanted with Met-1 tumor cells develop significantly more lung metastases than LFD-fed mice (Hillers-Ziemer et al., 2020). Similarly, HFD-fed mice had significantly more TC2 metastatic foci than LFD-fed mice (*p* = 0.03, Figure 1D). The metastases were variable in size, and diet did not significantly affect the sizes of metastatic foci. These results indicate that obesity also promotes pulmonary metastasis, in addition to accelerating mammary tumor growth.

Since obesity has been associated with promoting metastasis-initiating cells in breast cancer (Bousquenaud et al., 2018; Hillers-Ziemer et al., 2020), we examined the ability of tumor cells isolated from end-stage tumors from LFD and HFD-fed mice to establish metastases. Met-1 tumor cells were isolated primary tumors and injected tumor cells into the tail veins of recipient mice fed the LFD (Figure 1E). After 8 weeks, the lungs of transplanted mice exhibited no significant differences in the average number of metastatic foci, irrespective of the source of the tumor cells (Figure 1F). Together these data suggest that tumor extrinsic factors may contributed to metastasis under conditions of obesity.

### Obesity Enhances Myeloid Lineage Cells During Metastasis

To assess how obesity impacts the lungs to facilitate metastatic colonization, 3-week-old female FVB/N mice were fed the HFD or LFD for 16 weeks (Figure 2A), then Met-1 or TC2 tumor cells were injected into the tail vein to generate lung metastasis in the absence of a primary tumor. Metastases were given time to establish, then lung tissue was collected, and metastases were quantified in tissue sections. HFD-fed mice had significantly increased numbers of lung metastases than LFD-fed mice after injections of either Met-1 (*p* = 0.02, Figure 2B) or TC2 tumor cell lines (*p* = 0.04, Figure 2C). These results suggest that even in the absence of a primary tumor, obesity promotes metastatic colonization.

**Figure 2.**
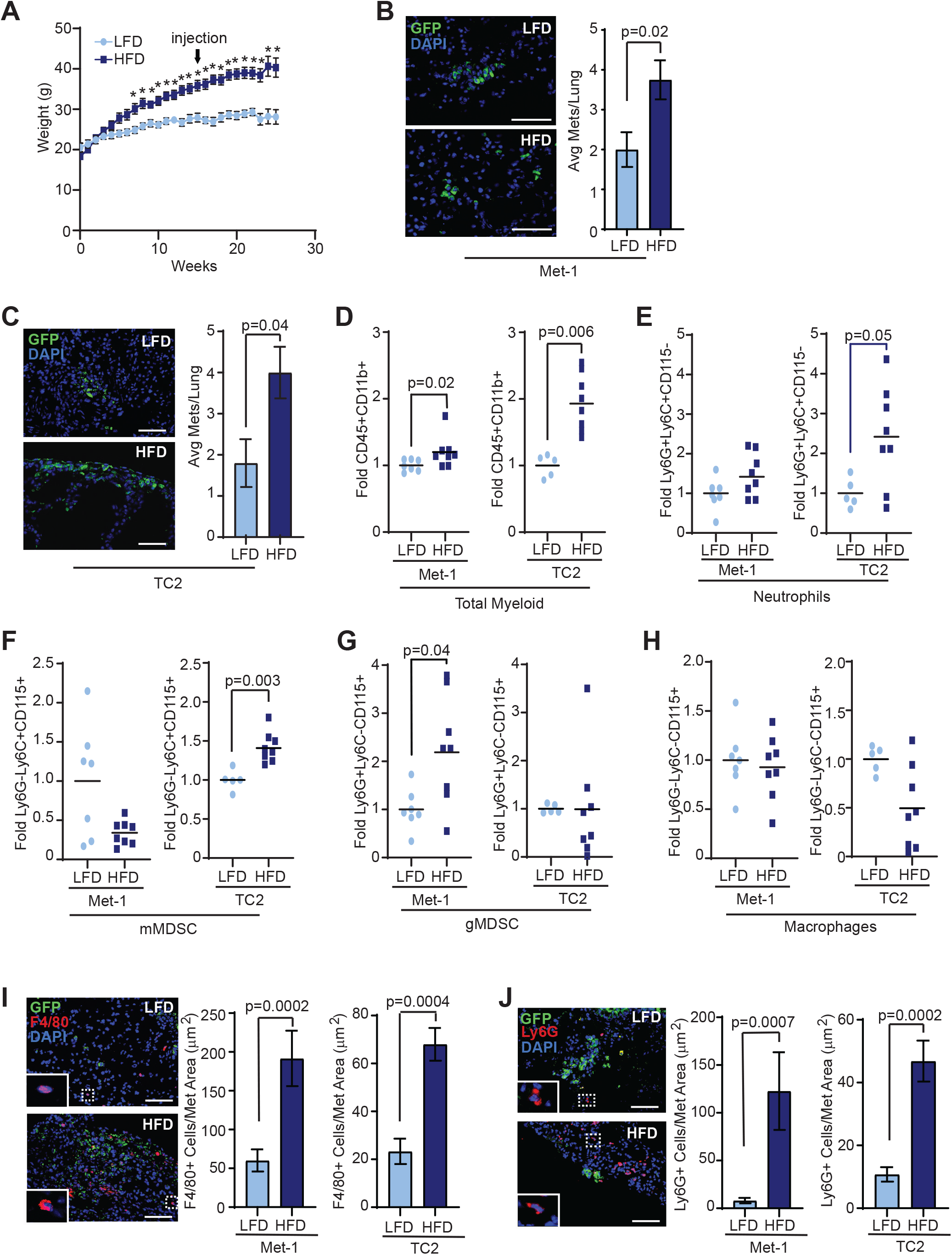
Obesity enhances myeloid lineage cells during metastasis. (A) Weight gain of mice fed LFD or HFD (n=12-17 mice/group). Arrow indicates the time point of tail vein injections of Met-1 or TC2 cells. Quantification of GFP^+^ metastatic foci in lungs of LFD and HFD-fed mice injected with Met-1 (B, n=7-9 mice/group) or TC2 (C, n=5-8 mice/group) tumor cells. Flow cytometry quantification of CD45^+^CD11b^+^ myeloid lineage cells (D), Ly6G^+^Ly6C^+^CD115^−^ neutrophils (E), Ly6G^−^Ly6C^+^CD115^+^ monocytic myeloid-derived suppressor cells (mMDSC, F), Ly6G^+^Ly6C^−^CD115^+^ granulocytic MDSC (gMDSC, G), and Ly6G^−^Ly6C^−^CD115^+^ macrophages (H) in lungs from mice injected with either Met-1 or TC2 tumor cells (n=5-8 mice/group). Results were normalized to CD45^+^CD11b^+^ cells and expressed as fold change compared to controls. (I) Quantification of F4/80^+^ macrophages surrounding lung metastases (n=5 mice/group). (J) Quantification of Ly6G^+^ neutrophils and gMDSC surrounding lung metastases. Magnification bars = 50μm.

Myeloid lineage cells help to promote tumor cell survival and growth at metastatic sites (Swierczak and Pollard, 2020). To determine the impact of obesity on myeloid lineage cells in pulmonary metastases, lungs from mice injected with tumor cells were dissociated into single cells, stained with antibodies for CD45, CD11b, Ly6G, Ly6C, and CD115, and myeloid lineage cell populations were analyzed using flow cytometry (Figure S1). In HFD-fed mice injected with either Met-1 or TC2 tumor cells, the total CD45^+^CD11b^+^ myeloid lineage cell population was significantly increased compared to LFD-fed mice (Figure 2D), indicating that obesity enhances myeloid cell recruitment during metastatic outgrowth. However, changes in specific myeloid lineage subpopulations differed with respect to the parental tumor cell line. While no significant difference in neutrophils were observed between LFD and HFD-fed mice bearing Met-1 metastases, HFD-fed mice with TC2 metastases had significantly increased Ly6C^+^Ly6G^+^CD115^−^ neutrophils compared to LFD-fed mice (*p* = 0.05, Figure 2E). Similarly, the population of Ly6C^+^Ly6G^+^CD115^+^ mMDSCs was comparable between HFD and LFD-fed Met-1 metastases-bearing mice, while mMDSCs were significantly elevated in HFD-fed mice with TC2 metastases compared to LFD-fed mice (*p* = 0.003, Figure 2F). Further, the population of Ly6C^−^Ly6G^+^CD115^+^ gMDSCs was significantly increased in HFD-fed mice with Met-1 metastases compared to those from LFD-fed mice (*p* = 0.04, Figure 2G), however there was no observed difference in gMDSCs in lungs from TC2 injected HFD or LFD-fed mice (Figure 2G). In contrast, Ly6C^−^Ly6G^−^CD115^+^ macrophages were unaltered between LFD and HFD-fed mice injected with either Met-1 or TC2 tumor cells (Figures 2H). Together, these data suggest that while obesity promotes myeloid lineage cell recruitment into lung tissue during metastasis, enrichment for specific immune cell types may depend upon properties of the tumor cells within the metastases.

To assess spatial relationships between myeloid lineage cells and pulmonary metastases, we examined populations of myeloid lineage cells using immunofluorescence. Since CD115 is expressed on numerous cell types including macrophages, mMDSCs, and gMDSCs (Hey et al., 2015), we utilized F4/80 expression to detect macrophages, which is expressed on both interstitial and alveolar lung macrophages (Zaynagetdinov et al., 2013). Tissue surrounding Met-1 and TC2 metastases in lungs of HFD-fed mice demonstrated significantly greater F4/80^+^ macrophage recruitment than metastases from LFD-fed mice (Figure 2I). Further, Ly6G^+^ cells, including both gMDSCs and neutrophils, were increased around metastases from HFD-fed mice compared to metastases from LFD-fed mice (Figure 2J). These data indicate that recruitment of myeloid lineage cells, including gMDSC and mMDSC, to metastatic sites in the lungs is enhanced by obesity.

### Obesity Alters the Complement of Myeloid Lineage Cells in the Lungs Prior to Metastasis

In the bone marrow, obesity enhances myeloid progenitor cell proliferation and upregulates cytokine production (Nagareddy et al., 2014; Singer et al., 2014), suggesting that immune cells are systemically increased as a result of obesity. We hypothesized that obesity may promote trafficking of myeloid lineage cells into the lungs prior to onset of primary tumors, which may enhance metastasis. To examine myeloid lineage cell recruitment into lungs prior to metastasis formation, we collected lungs from LFD and HFD-fed tumor-naïve mice (Figure 3A) and performed flow cytometry to quantify myeloid lineage cell populations. In contrast to metastatic lungs, there was no significant difference in total CD45^+^CD11b^+^ myeloid lineage cells among lungs from LFD and HFD-fed mice (Figure 3B). However, Ly6C^−^Ly6G^−^CD115^+^ macrophages (*p* = 0.01, Figure 3C) and Ly6C^−^Ly6G^+^CD115^+^ gMDSCs (*p* = 0.03, Figure 3D) were significantly increased while Ly6C^+^Ly6G^+^CD115^−^ neutrophils and Ly6C^+^Ly6G^+^CD115^+^ mMDSCs were not significantly different in lungs of HFD-fed mice compared those from LFD-fed mice (Figures 3E, F). These data indicate that obesity promotes recruitment of macrophages and gMDSCs to the lungs prior to metastasis formation.

**Figure 3.**
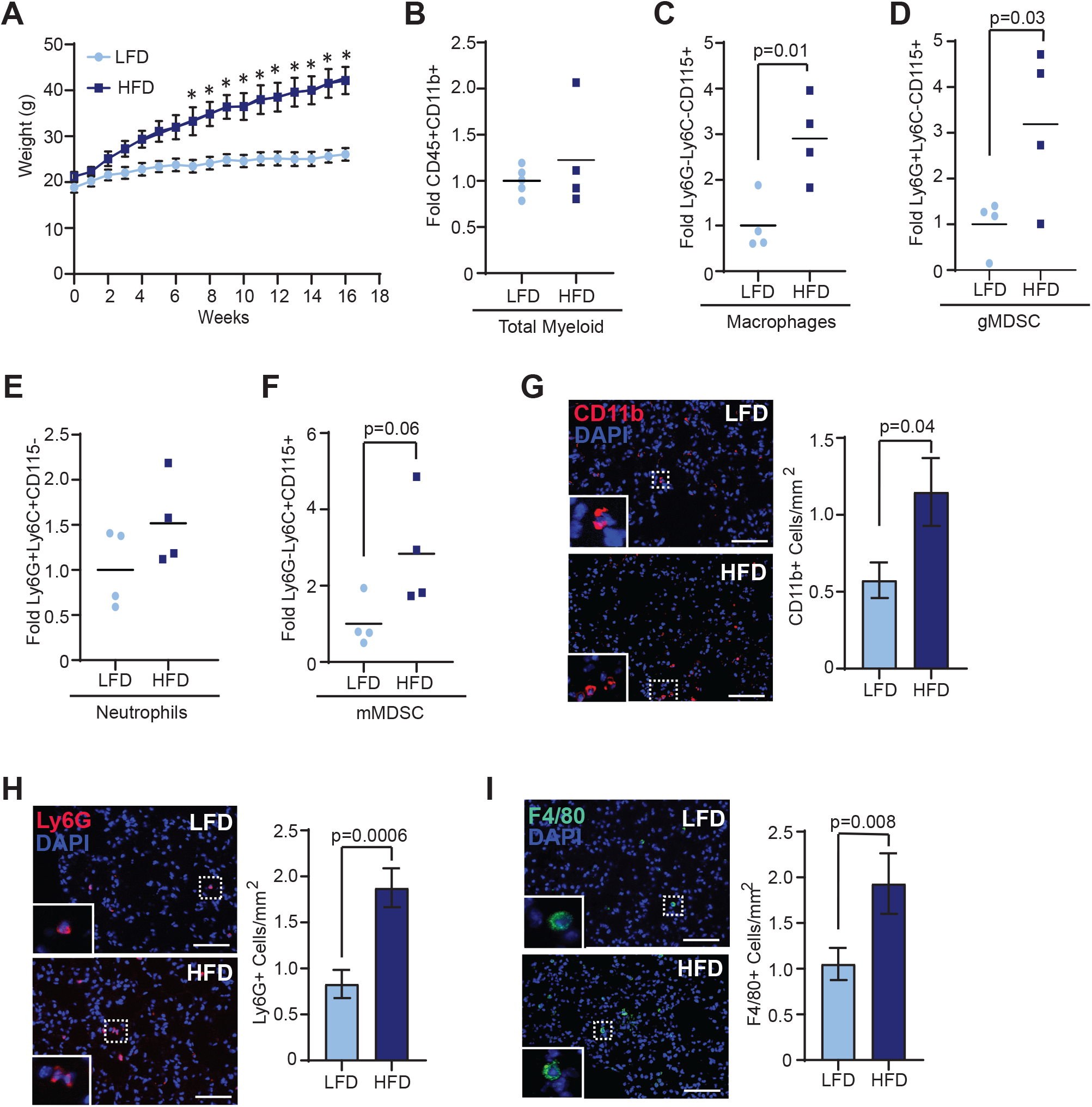
Obesity alters the compliment of myeloid lineage cells in lungs prior to metastasis. (A) Weight gain of female mice fed either LFD or HFD for 16 weeks (n=8 mice/group). Flow cytometry quantification of CD45^+^CD11b^+^ myeloid lineage cells (B), Ly6G^−^Ly6C^−^CD115^+^ macrophages (C), Ly6G^+^Ly6C^−^CD115^+^ gMDSC (D), Ly6G^+^Ly6C^+^CD115^−^ neutrophils (E), and Ly6G^−^Ly6C^+^CD115^+^ mMDSC (F). Results were normalized to CD45^+^CD11b^+^ cells and expressed as fold change compared to controls (n=4 mice/group). Quantification of CD11b^+^ myeloid lineage cells (G), Ly6G^+^ neutrophils and gMDSCs (H), and F4/80^+^ macrophages (I) in lungs of LFD and HFD-fed mice (n=5 mice/group). Magnification bars = 50μm.

To assess the localization of myeloid lineage cells in tumor-naïve mice, we examined immune cell markers in lungs sections from LFD and HFD-fed mice using immunofluorescence. HFD-fed mice demonstrated significantly increased numbers of CD11b^+^ myeloid lineage cells per area of lung tissue than LFD-fed mice (*p* = 0.03, Figure 3G). Further, recruitment of Ly6G^+^ gMDSCs and neutrophils (*p* = 0.0006, Figure 3H) and F4/80^+^ macrophages (*p* = 0.008, Figure 3I) were significantly increased in lung tissue of HFD-fed mice compared to LFD-fed mice. These data suggest that obesity alters myeloid lineage cell trafficking into the lungs before tumor formation.

### Obesity Activates Stromal Cells within the Lungs through TGFβ1 Expression

In tumor-bearing mice, pre-metastatic niches in the lungs have been shown to promote recruitment of bone marrow-derived immune cells through increased collagen deposition and fibrosis (Wong et al., 2011). To determine how obesity impacts collagen deposition within the lung microenvironment, we quantified collagen in the lungs of tumor-naïve LFD and HFD-fed mice using picrosirius red staining. The number of collagen fibers within lung tissue was significantly increased in HFD-fed mice compared to lungs from LFD-fed mice (*p* = 0.03, Figure 4A). Collagen fiber length and width remained unchanged between lungs of LFD and HFD-fed mice (Figure S2A). These results suggest that obesity increases accumulation of collagen fibers within lung tissue of tumor-naïve mice.

**Figure 4.**
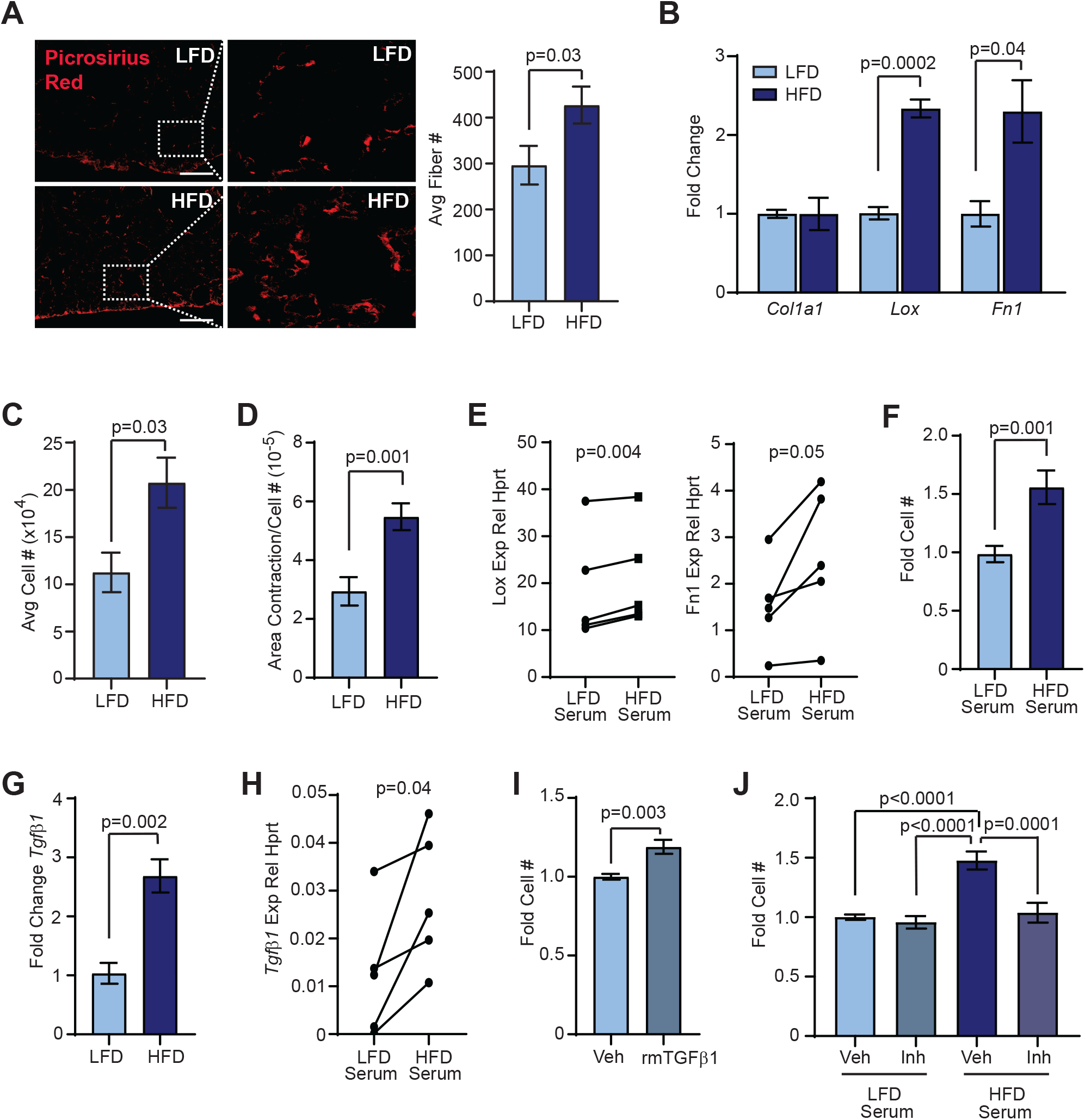
Obesity alters lung stromal cells within the lungs in the absence of tumor formation. (A) Quantification of picrosirius red-stained collagen fibers in lung sections of LFD and HFD-fed non-tumor-bearing mice (n=5 mice/group). (B) Expression of ECM components from primary lung stromal cells. Expression differences were normalized to *Hprt* and represented as fold change from controls (n=6-7 mice/group). (C) Cell numbers of isolated lung stromal cells grown in culture for 7 days (n=3 mice/group). (D) Average area of collagen contracted from lung stromal cells (n=3 mice/group). (E) *Lox* and *Fn1* expression in lung stromal cells isolated from LFD-fed mice treated with serum from LFD or HFD-fed mice. Represented as relative expression compared to *Hprt* (n=5 mice/group; paired t-test). (F) Cell numbers of isolated lung stromal cells from LFD-fed mice treated with serum from LFD or HFD-fed mice for 6 days (n=5 mice/group). (G) Expression of *Tgfβ1* from primary lung stromal cells. Expression differences were normalized to *Hprt* and represented as fold change from controls (n=6-7 mice/group). (H) *Tgfβ1* expression in lung stromal cells isolated from LFD-fed mice treated with serum from LFD or HFD-fed mice. Represented as relative expression compared to *Hprt* (n=5 mice/group; paired t-test). (I) Lung stromal cells from LFD-fed mice treated with vehicle (veh) or recombinant mouse (rm) TGFβ1 (n=3 mice/group). (J) Lung stromal cells from LFD-fed mice treated with serum from LFD or HFD-fed mice supplemented with veh or TGFβ inhibitor (inh) SB431542 (n=3 mice/group). Magnification bars = 50μm.

To examine how obesity may impact lung stromal cells, stromal cells were isolated from lung tissue of LFD and HFD-fed mice and cultured to generate adherent cells. These adherent lung stromal cell cultures did not contain detectable transcripts for *Cnn1*, *Cd31*, and *Epcam*, and only low *Cd45* expression compared to splenic tissue (Figure S2B), indicating that short-term culture of lung stromal cells depletes epithelial, endothelial, pericyte, and immune cell populations. Although lungs of HFD-fed mice exhibited higher numbers of collagen fibers, lung stromal cells from HFD-fed mice expressed similar levels of *Col1a1* compared to those from LFD-fed mice (Figure 4B). However, expression of *Lox* (lysyl oxidase), an enzyme that increases ECM crosslinking and collagen stability, was significantly increased in lung stromal cells from HFD-fed mice compared to LFD-fed mice (*p* = 0.0002, Figure 4B). Additionally, lung stromal cells from HFD-fed mice demonstrated significantly increased expression of *Fn1* (fibronectin; *p* = 0.04, Figure 4B), which has been implicated as an ECM component of cancer-induced pre-metastatic niches (Kaplan et al., 2005).

In culture, lung stromal cells from HFD-fed mice demonstrated significantly increased cell numbers after 7 days compared to those from LFD-fed mice, suggesting increased proliferation of stromal cells from HFD-fed mice (*p* = 0.03, Figure 4C). To test how obesity impacted lung stromal cell function, lung stromal cells were plated into collagen gels and contractility of the gel was measured after 7 days. Lung stromal cells from HFD-fed mice demonstrated significantly increased contraction of collagen gels compared to lung stromal cells from LFD-fed mice (*p*= 0.001, Figure 4D). Together, these results indicate that obesity alters ECM deposition and function of lung stromal cells.

In obesity, adipose tissue is chronically inflamed, and multiple inflammatory cytokines and growth factors are elevated systemically in serum (Ecker et al., 2019; Quail et al., 2017). We hypothesized that circulating inflammatory factors may promote the changes we observed in lung stromal cells from HFD-fed mice. To test this hypothesis, we cultured lungs stromal cells from LFD-fed mice and treated them with serum isolated from either LFD or HFD-fed mice. Consistent with our *in vitro* analyses of lung stromal cells isolated from HFD-fed mice, lung stromal cells from LFD-fed mice that were treated with serum from HFD-fed mice demonstrated significantly increased expression of *Lox* (*p* = 0.004) and *Fn1* (*p* = 0.05) compared to the same cells treated with serum from LFD-fed mice (Figure 4E). Lung stromal cells from LFD-fed mice treated with serum from HFD-fed mice also grew more rapidly, compared to the paired lung stromal cells treated with serum from LFD-fed mice (*p* = 0.001; Figure 4F), demonstrating that exposure to serum from HFD-fed mice enhanced lung stromal cell proliferation rates. Together, these results suggest that exposure to circulating inflammatory cytokines and/or growth factors from obese mice promotes expression of ECM remodeling components as well as functional changes of lung stromal cells.

Transforming growth factor beta (TGFβ) has been implicated in increased ECM production in pathological conditions of lung fibrosis, with TGFβ1 as the predominant TGFβ isoform expressed (Yue et al., 2010). Lung stromal cells from HFD-fed mice expressed significantly higher levels of *Tgfβ1* compared to those from LFD-fed mice (*p* = 0.002, Figure 4G). While TGFβ1 levels were similar in serum from LFD and HFD-fed mice (Figure S2C), treatment of lung stromal cells from LFD-fed mice with serum from HFD-fed mice resulted in significantly increased *Tgfβ1* expression (*p* = 0.04, Figure 4H). Functionally, treatment of lung stromal cells from LFD-fed mice with recombinant mouse TGFβ1 significantly enhanced proliferation (*p* = 0.003, Figure 4I). Further, treatment of lung stromal cells with serum from HFD-fed mice in the presence of TGFβ inhibitor SB431542 resulted in significantly reduced proliferation compared to serum from HFD-fed mice with vehicle (*p* = 0.0001, Figure 4J). In contrast, no differences were observed in proliferation of lung stromal cells from LFD-fed mice treated with serum from LFD-fed mice supplemented with vehicle or TGFβ inhibitor (Figure 4J). Together, these results indicate that inflammatory mediators from obese mice increase TGFβ1 expression within lung stromal cells to promote proliferation and collagen and ECM deposition.

### Obesity-Activated Lung Stromal Cells Enhance Migration of Bone Marrow and Tumor Cells

To assess how obesity-induced changes in lung stromal cells may aid in trafficking of bone marrow cells into the lungs, we tested the ability of bone marrow cells isolated from LFD-fed mice to migrate toward secreted factors from lung stromal cells of LFD and HFD-fed mice through collagen-coated transwells. We collected conditioned media from lung stromal cells, and we examined the ability of isolated bone marrow cells to migrate in response to conditioned media. Immune cells adherent to the bottom surface of the membranes were significantly increased in response to conditioned media from lung stromal cells from HFD-fed mice compared to LFD-fed mice (*p* = 0.02, Figure 5A). We also observed CD45^+^ bone marrow cells that invaded through the collagen into the conditioned media (Figure 5B). Approximately 90% of these cells expressed marker CD11b, consistent with cells of the myeloid lineage, and about 75% of the cells also expressed Ly6G (Figure 5B). Although conditioned media from HFD-fed mice did not significantly alter the types of cells that invaded through the collagen, the number of cells that were present in the conditioned media of lung stromal cells from HFD-fed mice was significantly increased compared to controls (*p* = 0.0007, Figure 5B). These data suggest that obesity-altered lung stromal cells enhance trafficking of bone marrow-derived immune cells into the lungs.

**Figure 5.**
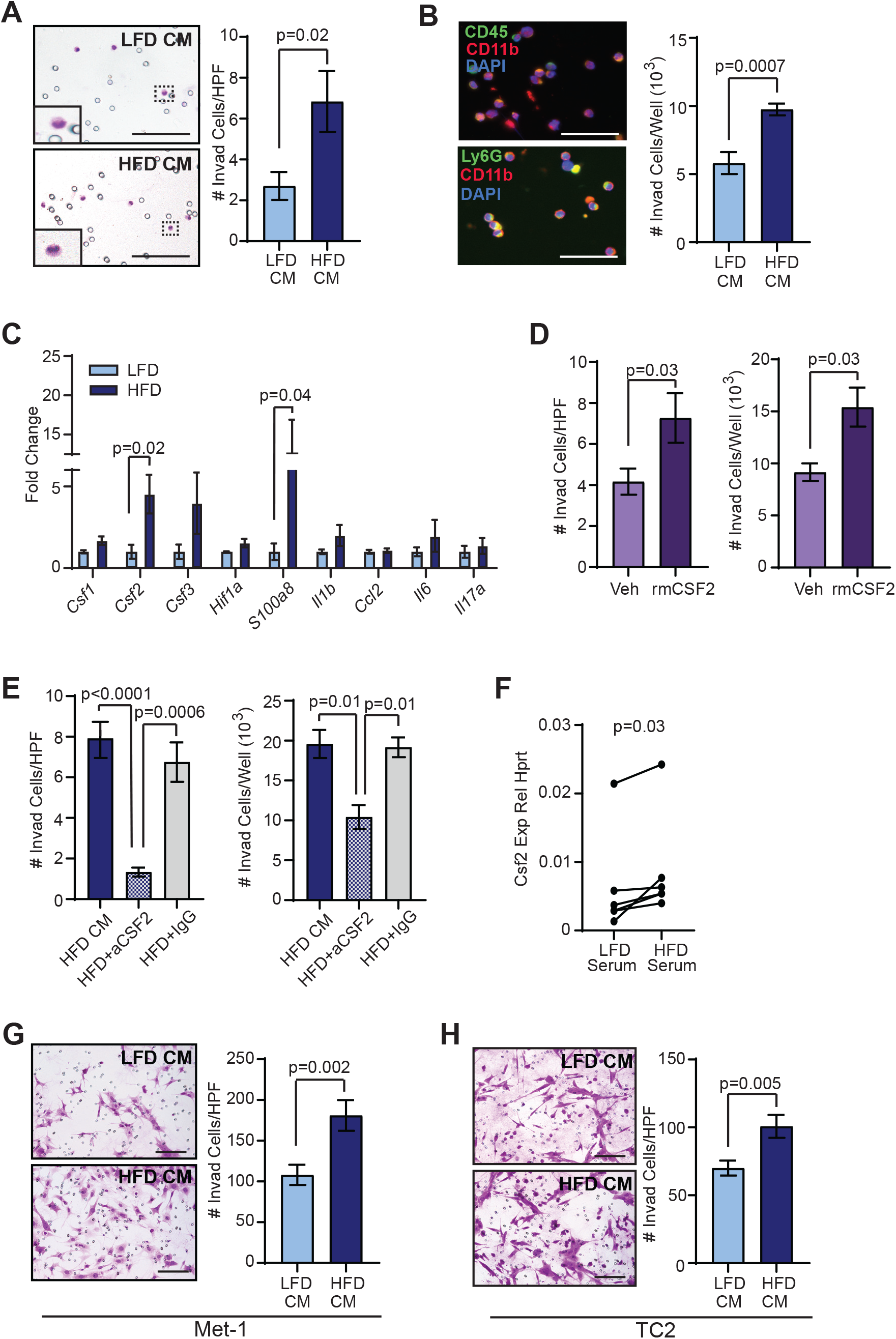
Lung stromal cells from HFD-fed mice recruit myeloid lineage cells through CSF2 expression. (A) Quantification of adherent bone marrow-derived cells invading toward conditioned media from lung stromal cells from LFD and HFD-fed mice (n=5 mice/group). (B) Quantification of non-adherent bone marrow-derived cells in conditioned media from lung stromal cells (n=5 mice/group). (C) Expression levels of cytokines from primary lung stromal cells. Differences were normalized to *Hprt* and represented as fold change from controls (n=6-7 mice/group). (D) Quantification of adherent and non-adherent bone marrow-derived cells invading toward vehicle or recombinant mouse CSF2 (n=3 experiments). (E) Quantification of adherent and non-adherent bone marrow-derived cells invading in response to conditioned media isolated from lung stromal cells from HFD-fed mice supplemented with vehicle, IgG antibodies, or blocking antibodies for CSF2. (F) *Csf2* expression in lung stromal cells isolated from LFD-fed mice treated with serum from LFD or HFD-fed mice. Represented as relative expression compared to *Hprt* (n=5 mice/group; paired t-test). (G) Quantification of Met-1 cells invading toward conditioned media from lung stromal cells (n=5 mice/group). (H) Quantification of TC2 cells invading toward conditioned media from lung stromal cells (n=5 mice/group). Magnification bars = 50μm.

Stromal cells in pre-metastatic secondary organs exhibit altered production and secretion of cytokines in response to tumor-derived factors (Liu and Cao, 2016), as metastatic tumor cells are reliant on local stromal cells to successfully colonize the organ (Lambert et al., 2017). Lung stromal cells from HFD-fed mice demonstrated significantly increased expression of *Csf2* (*p* = 0.02), which is elevated in local sites of inflammation and is implicated in myeloid lineage cell recruitment and maturation (Becher et al., 2016), and *S100a8* (*p* = 0.04), which modulates inflammatory responses through immune cell recruitment into tissues (Wang et al., 2018) (Figure 5C). Media containing recombinant mouse CSF2 significantly increased invasion of immune cells through collagen-coated transwells as either adherent cells (*p* = 0.03) or into the media (*p* = 0.03, Figure 5D). In addition, supplementation of conditioned media from lung stromal cells of HFD-fed mice with blocking antibodies for CSF2 significantly reduced invasion through transwells both for adherent cells (*p* = 0.0006) as well as cells within the media containing the blocking antibodies (*p* = 0.01) as compared to conditioned media from lung stromal cells of HFD-fed mice treated with IgG control antibodies (Figure 5E). Together, these results suggest that lung stromal cells from obese mice enhance myeloid lineage cell recruitment through elevated expression of CSF2.

Since lung stromal cells from LFD-fed mice were functionally altered by exposure to serum from HFD-fed mice, we hypothesized that serum collected from HFD-fed mice may induce *Csf2* expression in lung stromal cells from LFD-fed mice. As shown in Figure 5F, lung stromal cells from LFD-fed mice significantly increased expression of *Csf2* in response to treatment with serum from HFD-fed mice, compared to serum from LFD-fed mice (*p* = 0.03). Together, these results suggest that circulating factors induced by obesity can promote *Csf2* expression in lung stromal cells.

During lung metastasis, disseminated tumor cells leave blood vessels and invade into the lung stroma. We hypothesized that activated lung stromal cells from obese mice may also promote the invasion of tumor cells. Similar to our observations of immune cells, increased numbers of Met-1 (*p* = 0.002; Figure 5G) and TC2 (*p* = 0.005; Figure 5H) tumor cells invaded through collagen-coated transwells in response to factors secreted by lung stromal cells from HFD-fed mice. These results suggest that obesity enhances recruitment of immune cells and tumor cells together into the lung stroma through activation of lung stromal cells.

## DISCUSSION

Prior to metastasis, primary tumors create niches in distal organs conducive to metastatic colonization. In the absence of a primary mammary tumor, we observed that obesity altered the microenvironment of the lungs with similarities to tumor-induced pre-metastatic niches. Within lung tissue, obesity enhanced recruitment of myeloid lineage cells, in particular macrophages and gMDSC, as well as increased collagen fibers. Lung stromal cells isolated from obese mice demonstrated elevated expression of *Lox* and *Fn1,* similar to lung fibroblasts within pre-metastatic niches (Pein et al., 2020). These changes in the immune microenvironment and ECM under conditions of obesity enhanced the ability of ERα^+^ and ERα^−^ tumor cells to establish metastatic colonization. Within the mammary gland, obesity also promoted the rapid growth of ERα^+^ and ERα^−^ primary mammary tumors, and obesity has been shown to enhance cancer stem-like cells within mammary tumors (Bowers et al., 2018; Hillers-Ziemer et al., 2020). Since cancer stem-like cells may increase metastatic potential (Baccelli and Trumpp, 2012), these results suggest that obesity enhances lung metastasis both through promotion of tumor cells with increased metastatic potential as well as by establishment of favorable conditions for metastasis. These obesity-induced conditions may lead to the clinically observed increased risk for metastasis in obese breast cancer patients (Ewertz et al., 2011; Sestak et al., 2010).

Similar to cancer, obesity induces non-resolving inflammation, resulting in enhanced circulating numbers of bone marrow-derived myeloid lineage cells (Nagareddy et al., 2014; Singer et al., 2014). Our results indicated that these bone marrow-derived cells may be recruited from the circulation into lung tissue by lung stromal cells in obesity through locally elevated CSF2 as well as S100A8, which has been shown to promote accumulation of myeloid lineage cells at metastatic sites *in vivo* (Hiratsuka et al., 2006; Hiratsuka et al., 2008). Recent studies also suggest that IL-5 expression from obese adipose tissue may also enhance homing of myeloid lineage cells, in particular neutrophils, to the lungs (Quail et al., 2017). In pre-metastatic niches, recruited myeloid lineage cells prepare distant organs for metastatic seeding (Liu and Cao, 2016; Swierczak and Pollard, 2020). Macrophages are necessary for metastatic growth in the lungs (Linde et al., 2018; Qian et al., 2009; Qian et al., 2015), and obesity may alter macrophage function in lung tissue, potentially in response to circulating inflammatory cytokines (Manicone et al., 2016). gMDSCs have also been shown to enhance metastasis through immunosuppressive effects and vascular remodeling within the lungs (Yan et al., 2010). Our studies support a role for increased early recruitment of myeloid lineage cells into lung tissue which may further promote an environment conducive to metastasis under conditions of obesity.

ECM remodeling in distant organs is essential for metastatic colonization (Sleeman, 2012). We observed that obesity enhances collagen deposition and *Lox* expression within lung tissue of tumor-naïve mice. Collagen deposition and stabilization in the lungs through activity of lysyl oxidase has been shown to promote breast cancer cell metastatic colonization (Elia et al., 2019). Lung stromal cells from obese tumor-naïve mice also exhibited increased proliferation rates and enhanced contractility, which has been observed in lung fibroblasts isolated from lung metastatic sites (Pein et al., 2020). Lung stromal cells from obese mice expressed significantly higher levels of *Tgfβ1*, and treatment with serum from obese mice induced *Tgfβ1* expression in lung stromal cells from LFD-fed mice. In models of lung fibrosis, enhanced TGFβ1 expression precedes increased collagen and extracellular matrix deposition (Hoyt and Lazo, 1988; Yi et al., 1996). These changes in lung fibroblasts from obese mice may be due to exposure to inflammatory cytokines present in serum, as treatment with tumor-derived factors in culture resulted in increased expression of collagen-1A1 and myofibroblast marker smooth muscle actin in lung fibroblasts (Kong et al., 2019). Leptin, which is an adipokine that is increased in the serum of obese individuals (Maffei et al., 1995), may also play a role (Watanabe et al., 2019). In pulmonary fibrosis, lung fibroblasts are activated through increases in microRNAs (Liu et al., 2010; Souma et al., 2018), immune cells (Li et al., 2018), and serum cytokines (Su et al., 2016), and further studies are needed to determine the mediators of obesity-induced lung stromal activation.

Metastasis is the primary cause of breast cancer mortality and identifying points of intervention to reduce metastatic risk in obese breast cancer patients is critical. Given that obesity is a chronic inflammatory disease, targeting obesity-induced recruitment of immune cells to the lungs may have therapeutic benefits for obese breast cancer patients. A recent study demonstrated that 10% loss of body mass in a small cohort of morbidly obese individuals resulted in a decrease in systemic inflammatory markers (Alemán et al., 2017), which suggests that weight loss may reduce systemic inflammation. However, other studies have suggested that weight loss may not reverse epigenetic changes induced in obese adipose tissue (Rossi et al., 2016), and the impact of weight loss on the function of lung stromal cells needs to be investigated. Given the similarities that we observed among lung stromal cells in obesity and cancer-associated fibroblasts, therapeutics in development to target inflammatory characteristics of fibroblasts as well as anti-fibrotic agents may also have efficacy to limit metastasis in obese breast cancer patients (Chen, 2019; Liu et al., 2019). Further studies are necessary to determine how obesity alters other frequent sites of breast cancer metastases. Given that obesity increases the incidence of multiple types of cancer (Bhaskaran et al., 2014), understanding how obesity promotes early metastasis may improve treatment options for the rising population of obese cancer patients.

## ACKNOWLEDGEMENTS

The authors would like to thank Brenna Walton and Genevra Kuziel for helpful discussions, and the University of Wisconsin Carbone Cancer Flow Cytometry Core for technical expertise. This work was supported by NIH grants R01 CA227542 (to L.M.A.), T32 GM007215 (to L.E.H.), R25 GM083252 (to A.W.), T35 OD011078 (to A.S.) and University of Wisconsin Carbone Cancer Center Support Grant P30 CA014520.

## AUTHOR CONTRIBUTIONS

L.E.H. and L.M.A. initiated the study, performed analyses and prepared figures. L.E.H., A.J., A.S., C.G., and A.W. performed immunofluorescence and analyses, V.T. and L.E.H. performed qRT-PCR experiments and in vitro studies and analyses, L.E.H., A.W. and L.M.A. provided input for research design and interpretation and edited the manuscript, L.M.A. supervised the study, and L.E.H., A.W. and L.M.A. wrote and revised the manuscript.

## METHODS AND MATERIALS

### Animal Studies

All procedures involving animals were approved by the University of Wisconsin-Madison Institutional Animal Care and Use Committee (Animal Welfare Assurance Number (D16-00239)). Female FVB/NTac mice were purchased from Taconic Laboratories and maintained according to the Guide for Care and Use of Laboratory Animals in AAALAC-accredited facilities. Three-week-old FVB/N female mice were randomized to be fed low-fat diet (LFD, 10% kcal from fat, Test Diet; 58Y2) or high-fat diet (HFD, 60% kcal from fat, Test Diet; 58Y1) for 16 weeks to induce obesity. Purified diets contained equal amounts of vitamins and micronutrients. Body mass was measured weekly.

### Cell Lines

Met-1 cells were provided by Dr. Alexander Borowsky (Borowsky et al., 2005) and were transduced with lentivirus encoding green fluorescent protein (GFP) as described (Hillers-Ziemer et al., 2020). TC2 GFP^+^ cells were provided by Dr. Linda Schuler (Barcus et al., 2017). Primary lung stromal cells were isolated from lungs of LFD and HFD-fed mice. Lung tissue was digested for 1 hr in DMEM:F12 (Corning; 10-090-CV) supplemented with 3 mg/mL collagenase I (MilliporeSigma; 1148089). Digested tissue was incubated for 2 hr to collect adherent cells, and adherent cells were expanded in culture for no more than three passages prior to use in assays. Met-1 tumor cells were cultured in DMEM (Corning; 10-017-CV) supplemented with 10% FBS, lung stromal cells were cultured in DMEM supplemented with 10% calf serum, and TC2 cells were cultured in DMEM supplemented with 10% FBS and 1 mg/mL G418 (ThermoFisher Scientific; 11811023). All media contained 1% antibiotic/anti-mycotic solution, and cells were maintained at 37°C in 5% CO_2_. Tumor cell lines were not validated and were tested for mycoplasma prior to use in experiments (Idexx Bioresearch).

### Tumor Cell Transplantations

To generate tumors, 5×10^5^ Met-1 or 2.5×10^4^ TC2 cells were suspended in 2:1 Matrigel (Corning; 354234):DMEM and injected bilaterally into the inguinal mammary glands of LFD or HFD-fed FVB/N female mice. Tumor diameters were measured three times each week using calipers. Tumor volume was calculated using the formula 4/3πr^3^. To generate metastases, 5×10^5^ Met-1 or TC2 cells were suspended in sterile PBS and injected into the tail vein of HFD or LFD-fed mice. End stage for metastatic development was defined as 6-weeks post tail vein injections for mice transplanted with Met-1 cells or 8-weeks for TC2 recipient mice.

### Conditioned Media and Invasion Assays

Lung stromal cells were grown on 100-mm plates until confluent. Cells were washed with PBS, then grown for 24 hr in DMEM supplemented with 0.5% calf serum and 1% antibiotic/antimycotic solution. Conditioned media was filtered through 0.22-μm filters (ThermoFisher Scientific; 09-720-004). To assess invasion, 1×10^5^ bone marrow cells from LFD-fed mice were plated in duplicate in serum-free media on inserts with 8 μm pores (Corning; 353097) coated with 1 mg/mL Type I rat tail collagen (Corning; 354236), and invasion toward conditioned media from lung stromal cells was measured after 4 hr. Inserts were formalin-fixed and stained with 0.1% crystal violet. Four images of each invasion insert were taken at 100x magnification on a Nikon Eclipse E600 Microscope with a QICAM Fast 1394 camera and quantified using ImageJ (NIH) with cell counter plug-in. Bone marrow cells that invaded into the conditioned media were quantified using a hemocytometer, then cytospun onto slides, fixed in methanol and stained using antibodies for CD45 (ThermoFisher Scientific; 14-0451-82), CD11b (Novus Biologicals; NB110-89474), Ly6G (Abcam; ab25377). Invasion was also quantified in response to serum-free DMEM supplemented with 10 ng/mL of recombinant mouse CSF2 (R&D Systems, 415-ML-5), PBS or conditioned media from lung stromal cells from HFD-fed mice treated with 5 μg/mL of either CSF2 neutralizing antibodies (R&D Systems, MAB415-100) or rat IgG antibodies (R&D Systems, 6-001-A) for 1 hr prior to the assay start. Four biological replicates were tested for each condition.

### Proliferation Assays

To quantify differences in proliferation, 1×10^5^ lung stromal cells were plated in DMEM supplemented with 10% calf serum and 1% antibiotic/antimycotic solution. For serum treatment experiments, 1×10^5^ lung stromal cells from LFD-fed mice were plated with DMEM+5% serum collected from LFD or HFD-fed mice+1% antibiotic/antimycotic solution. To test responses to TGFβ1, lung stromal cells from LFD-fed mice were grown in 5% serum collected from LFD or HFD-fed mice supplemented with 10 μM of TGFβ inhibitor SB431542 (A10826A, Adooq Biosciences) or DMSO control or in DMEM +0.5% calf serum supplemented with 5 ng/mL recombinant mouse TGFβ1 (7666-MB, Bio-Techne Corporation) or PBS vehicle control. Cells were fed with media supplemented with serum every 2 days. All proliferation assays were plated in triplicate and counted after 6 days with a hemocytometer, then pelleted for RNA extraction. Latent TGFβ1 was activated prior to quantification, and total TGFβ1 was measured in serum diluted 1:50 in buffer from LFD and HFD-fed mice using TGFβ1 DuoSet ELISA (DY1679, Bio-Techne Corporation) in duplicate according to the manufacturer’s instructions.

### Contractility Assays

Type I rat tail collagen (Corning, 354236) was diluted and neutralized in an equal volume of filter sterilized HEPES. 5×10^4^ lung stromal cells were added for a final concentration of 2 mg/mL collagen. The neutralized collagen and cell mixture were plated on 6-well plates in triplicate and incubated at 37°C with 4 biological replicates. After 4 hours, the gels were released and floated in 2 mL of DMEM supplemented with 10% calf serum and 1% antibiotic/antimycotic solution. The gel diameter was measured with a ruler at day 0, 2, 4, 5, and 7. Gels were fed after measurement on days 2 and 4. Contracted area was calculated using A = πr^2^ by subtracting the area measured on day 7 from day 0. On day 7, gels were digested with collagenase for 10 min at 37°C, and the difference in area of contraction was divided by the number of cells in the gel at day 7.

### Quantitative RT-PCR

RNA was isolated from lung stromal cells using TRIzol (Life Technologies; 15596026) and purified using Qiagen RNeasy Mini Kit (Qiagen; 74104). RNA was reverse transcribed using the High Capacity cDNA Reverse Transcription Kit (Applied Biosciences; 4368814) and Techne Thermal Cycler. Quantitative RT-PCR was performed using iTaq SYBR Green Supermix (Bio-Rad; 172-5121) with a Bio-Rad CFX Connect Real-Time PCR Detection System (Bio-Rad). Transcripts were normalized to housekeeping gene hypoxanthine-guanine phosphoribosyltransferase (Hprt) and data was analyzed using the ΔΔCq method (fold change) or ΔCq method (relative expression). Primer sequences are listed in Table S1.

### Flow Cytometry

Lung tissue was dissociated to single cells and resuspended at 1×10^6^ cells/mL in PBS containing 1% calf serum. Dissociated lung cells were blocked with CD16/32 antibodies (ThermoFisher Scientific; 14-0161-82) for 30 minutes at 4°C. Cells were stained with fixable viability dye eFluor 780 (ThermoFisher Scientific; 65-0865-14) per manufacturers’ instructions, then incubated with antibodies to detect CD45-PE (0.5 μg/μL; ThermoFisher Scientific; 12-0451-81), CD11b-APC (0.5 μg/μL; ThermoFisher Scientific; 17-0112-82), Ly6C-eFluor 700 (1.5 μg/μL; Biolegend; 128023), Ly6G-BV421 (1 μg/μL; Biolegend; 127627), and CD115-PE-Dazzle594 (1 μg/μL; Biolegend; 135527) for 30 min at 4°C. Antibody-bound cells were resuspended at 1×10^6^ cells/mL and quantified using an Attune Nxt Flow Cytometer (ThermoFisher Scientific). Flow cytometry data were analyzed with the FlowJo software package v 10 (FlowJo, LLC). Gates were set on fluorescence minus one controls and analyzed according to published guidelines (Alexander et al., 2009).

### Histology and Immunofluorescence

Paraffin-embedded metastatic and non-metastatic lungs were sectioned and stained with Hematoxylin and Eosin (H&E) by the Experimental Pathology Laboratory (Carbone Cancer Center, University of Wisconsin-Madison). Tissue staining for estrogen receptor alpha (ERα; Santa Cruz; sc-8005), F4/80 (Biolegend; 123102), CD11b (Novus Biologicals; NB110-89474), Ly6G (Abcam; ab25377), and GFP (Novus Biologicals; NB100-1678) was performed as previously described (Arendt et al., 2013). CD11b^+^, F4/80^+^, and Ly6G^+^ cells were quantified in non-metastatic and metastatic lungs using ImageJ (NIH) and divided by either the total lung tissue area on each image or by the metastatic area on the image. Blinded tissue sections were imaged using a Nikon Eclipse E600 Microscope and QICAM Fast 1394 camera. Five images were taken for each lung and quantified from 5-9 lungs/group. To quantify metastases, clusters of 5 or more GFP^+^ Met-1 or TC2 tumor cells were considered a metastatic lesion.

### Collagen Quantification

Paraffin embedded lung sections from LFD and HFD-fed mice were deparaffinized and rehydrated through alcohols. Slides were incubated for 1 hr in picrosirius red solution [0.5 g of Direct Red 80 (Sigma-Aldrich; 2610-10-8) in 500 mL of saturated picric acid (Sigma-Aldrich; P6744-1GA)]. Slides were washed twice with acidified water (0.5% acetic acid) for 10 minutes, dehydrated in graded ethanol and xylenes, and mounted using Richard-Allan mounting medium (ThermoFisher Scientific; 4112APG). Imaging of picrosirius red was performed using a Nikon Eclipse E600 Microscope and QICAM Fast 1394 camera. Collagen fluorescence was detected using a TRITC filter cube and images were taken at 200x magnification. To remove the autofluorescent background, images were captured using a FITC filter cube at 200x magnification. ImageJ Image Calculator was utilized to subtract background autofluorescence from collagen fibers. After the background was removed, images were converted to 8-bit. Collagen fiber length, width, and number were measured using CT-FIRE detection software (LOCI; Madison, WI) (Bredfeldt et al., 2014).

### Statistical Analysis

Results were expressed as mean ± SEM unless otherwise stated. Data were tested with the Kolmogorov-Smirnov test for normality. Unless stated in the figure legends, statistical differences were determined using Student’s t-test for comparison of two groups or Analysis of Variance (ANOVA) with Tukey’s Multiple Comparison post-test for multiple groups. Differences in tumor growth rates and body weight differences over time were detected using two-way ANOVA with Tukey’s post hoc test. For serum treatments, differences were detected using paired t-tests. Sample numbers (n) are included in the figure legends for each experiment. P-values of 0.05 or less were considered significant. Statistical analyses were conducted using GraphPad Prism 8.3.1 (GraphPad Software).

## REFERENCES

Alemán, J.O., Iyengar, N.M., Walker, J.M., Milne, G.L., Da Rosa, J.C., Liang, Y., Giri, D.D., Zhou, X.K., Pollak, M.N., Hudis, C.A., et al. (2017). Effects of rapid weight loss on systemic and adipose tissue inflammation and metabolism in obese postmenopausal women. J Endocr Soc 1, 625–637.

Alexander, C.M., Puchalski, J., Klos, K.S., Badders, N., Ailles, L., Kim, C.F., Dirks, P., and Smalley, M.J. (2009). Separating stem cells by flow cytometry: reducing variability for solid tissues. Cell Stem Cell 5, 579–583.

Arendt, L.M., McCready, J., Keller, P.J., Baker, D.D., Naber, S.P., Seewaldt, V., and Kuperwasser, C. (2013). Obesity promotes breast cancer by CCL2-mediated macrophage recruitment and angiogenesis. Cancer Res 73, 6080–6093.

Baccelli, I., and Trumpp, A. (2012). The evolving concept of cancer and metastasis stem cells. J Cell Biol 198, 281–293.

Barcus, C.E., O’Leary, K.A., Brockman, J.L., Rugowski, D.E., Liu, Y., Garcia, N., Yu, M., Keely, P.J., Eliceiri, K.W., and Schuler, L.A. (2017). Elevated collagen-I augments tumor progressive signals, intravasation and metastasis of prolactin-induced estrogen receptor alpha positive mammary tumor cells. Breast Cancer Res 19, 9.

Becher, B., Tugues, S., and Greter, M. (2016). GM-CSF: From growth factor to central mediator of tissue tnflammation. Immunity 45, 963–973.

Bhaskaran, K., Douglas, I., Forbes, H., dos-Santos-Silva, I., Leon, D.A., and Smeeth, L. (2014). Body-mass index and risk of 22 specific cancers: a population-based cohort study of 5·24 million UK adults. The Lancet 384, 755–765.

Borowsky, A.D., Namba, R., Young, L.J., Hunter, K.W., Hodgson, J.G., Tepper, C.G., McGoldrick, E.T., Muller, W.J., Cardiff, R.D., and Gregg, J.P. (2005). Syngeneic mouse mammary carcinoma cell lines: two closely related cell lines with divergent metastatic behavior. Clin Exp Metastasis 22, 47–59.

Bousquenaud, M., Fico, F., Solinas, G., Rüegg, C., and Santamaria-Martínez, A. (2018). Obesity promotes the expansion of metastasis-initiating cells in breast cancer. Breast Cancer Res 20, 104.

Bowers, L.W., Rossi, E.L., McDonell, S.B., Doerstling, S.S., Khatib, S.A., Lineberger, C.G., Albright, J.E., Tang, X., deGraffenried, L.A., and Hursting, S.D. (2018). Leptin signaling mediates obesity-associated CSC enrichment and EMT in preclinical TNBC models. Mol Cancer Res 15, 869–879.

Bredfeldt, J.S., Liu, Y., Pehlke, C.A., Conklin, M.W., Szulczewski, J.M., Inman, D.R., Keely, P.J., Nowak, R.D., Mackie, T.R., and Eliceiri, K.W. (2014). Computational segmentation of collagen fibers from second-harmonic generation images of breast cancer. J Biomed Opt 19, 16007.

Chamberlin, T., D’Amato, J.V., and Arendt, L.M. (2017). Obesity reversibly depletes the basal cell population and enhances mammary epithelial cell estrogen receptor alpha expression and progenitor activity. Breast Cancer Res 19, 128.

Chen, D. (2019). Dually efficacious medicine against fibrosis and cancer. Med Sci (Basel) 7, 41.

Chun, J., El-Tamer, M., Joseph, K.A., Ditkoff, B.A., and Schnabel, F. (2006). Predictors of breast cancer development in a high-risk population. Am J Surg 192, 474–477.

Clements, V.K., Long, T., Long, R., Figley, C., Smith, D.M.C., and Ostrand-Rosenberg, S. (2018). Frontline Science: High fat diet and leptin promote tumor progression by inducing myeloid-derived suppressor cells. J Leukoc Biol 103, 395–407.

Dao, M.C., Saltzman, E., Page, M., Reece, J., Mojtahed, T., Wu, D., and Meydani, S.N. (2020). Lack of differences in inflammation and T cell-mediated function between young and older women with obesity. Nutrients 12, 237.

Ecker, B.L., Lee, J.Y., Sterner, C.J., Solomon, A.C., Pant, D.K., Shen, F., Peraza, J., Vaught, L., Mahendra, S., Belka, G.K., et al. (2019). Impact of obesity on breast cancer recurrence and minimal residual disease. Breast Cancer Res 21, 41.

Elia, I., Rossi, M., Stegen, S., Broekaert, D., Doglioni, G., van Gorsel, M., Boon, R., Escalona-Noguero, C., Torrekens, S., Verfaillie, C., et al. (2019). Breast cancer cells rely on environmental pyruvate to shape the metastatic niche. Nature 568, 117–121.

Ewertz, M., Jensen, M.B., Gunnarsdottir, K.A., Hojris, I., Jakobsen, E.H., Nielsen, D., Stenbygaard, L.E., Tange, U.B., and Cold, S. (2011). Effect of obesity on prognosis after early-stage breast cancer. J Clin Oncol 29, 25–31.

Ferrante, A.W., Jr. (2013). The immune cells in adipose tissue. Diabetes Obes Metab 15 Suppl 3, 34–38.

Gabrilovich, D.I. (2017). Myeloid-Derived Suppressor Cells. Cancer Immunol Res 5, 3–8.

Hey, Y.Y., Tan, J.K., and O’Neill, H.C. (2015). Redefining myeloid cell subsets in murine spleen. Front Immunol 6, 652.

Hillers-Ziemer, L.E., McMahon, R.Q., Hietpas, M., Paderta, G., LeBeau, J., McCready, J., Arendt, L.M. (2020). Obesity promotes cooperation of cancer stem-like cells and macrophages to enhance mammary tumor angiogenesis. Cancers (Basel) 12, 502.

Hillers, L.E., D’Amato, J.V., Chamberlin, T., Paderta, G., and Arendt, L.M. (2018). Obesity-activated adipose-derived stromal cells promote breast cancer growth and invasion. Neoplasia 20, 1161–1174.

Hiratsuka, S., Watanabe, A., Aburatani, H., and Maru, Y. (2006). Tumour-mediated upregulation of chemoattractants and recruitment of myeloid cells predetermines lung metastasis. Nat Cell Biol 8, 1369–1375.

Hiratsuka, S., Watanabe, A., Sakurai, Y., Akashi-Takamura, S., Ishibashi, S., Miyake, K., Shibuya, M., Akira, S., Aburatani, H., and Maru, Y. (2008). The S100A8-serum amyloid A3-TLR4 paracrine cascade establishes a pre-metastatic phase. Nat Cell Biol 10, 1349–1355.

Hoyt, D.G., and Lazo, J.S. (1988). Alterations in pulmonary mRNA encoding procollagens, fibronectin and transforming growth factor-beta precede bleomycin-induced pulmonary fibrosis in mice. J Pharmacol Exp Ther 246, 765–771.

Ioannides, S.J., Barlow, P.L., Elwood, J.M., and Porter, D. (2014). Effect of obesity on aromatase inhibitor efficacy in postmenopausal, hormone receptor-positive breast cancer: a systematic review. Breast Cancer Res Treat 147, 237–248.

Kaplan, R.N., Riba, R.D., Zacharoulis, S., Bramley, A.H., Vincent, L., Costa, C., MacDonald, D.D., Jin, D.K., Shido, K., Kerns, S.A., et al. (2005). VEGFR1-positive haematopoietic bone marrow progenitors initiate the pre-metastatic niche. Nature 438, 820–827.

Karatas, F., Erdem, G.U., Sahin, S., Aytekin, A., Yuce, D., Sever, A.R., Babacan, T., Ates, O., Ozisik, Y., and Altundag, K. (2017). Obesity is an independent prognostic factor of decreased pathological complete response to neoadjuvant chemotherapy in breast cancer patients. Breast 32, 237–244.

Kong, J., Tian, H., Zhang, F., Zhang, Z., Li, J., Liu, X., Li, X., Liu, J., Li, X., Jin, D., et al. (2019). Extracellular vesicles of carcinoma-associated fibroblasts creates a pre-metastatic niche in the lung through activating fibroblasts. Mol Cancer 18, 175.

Lahmann, P.H., Hoffmann, K., Allen, N., van Gils, C.H., Khaw, K.T., Tehard, B., Berrino, F., Tjønneland, A., Bigaard, J., Olsen, A., et al. (2004). Body size and breast cancer risk: findings from the European Prospective Investigation into Cancer And Nutrition (EPIC). Int J Cancer 111, 762–771.

Lambert, A.W., Pattabiraman, D.R., and Weinberg, R.A. (2017). Emerging biological principles of metastasis. Cell 168, 670–691.

Lauby-Secretan, B., Scoccianti, C., Loomis, D., Grosse, Y., Bianchini, F., and Straif, K. (2016). Body fatness and cancer--Viewpoint of the IARC Working Group. N Engl J Med 375, 794–798.

Li, Y., Bao, J., Bian, Y., Erben, U., Wang, P., Song, K., Liu, S., Li, Z., Gao, Z., and Qin, Z. (2018). S100A4(+) Macrophages are necessary for pulmonary fibrosis by activating lung fibroblasts. Front Immunol 9, 1776.

Linde, N., Casanova-Acebes, M., Sosa, M.S., Mortha, A., Rahman, A., Farias, E., Harper, K., Tardio, E., Reyes Torres, I., Jones, J., et al. (2018). Macrophages orchestrate breast cancer early dissemination and metastasis. Nat Commun 9, 21.

Liu, G., Friggeri, A., Yang, Y., Milosevic, J., Ding, Q., Thannickal, V.J., Kaminski, N., and Abraham, E. (2010). miR-21 mediates fibrogenic activation of pulmonary fibroblasts and lung fibrosis. J Exp Med 207, 1589–1597.

Liu, T., Han, C., Wang, S., Fang, P., Ma, Z., Xu, L., and Yin, R. (2019). Cancer-associated fibroblasts: an emerging target of anti-cancer immunotherapy. J Hematol Oncol 12, 86.

Liu, Y., and Cao, X. (2016). Characteristics and significance of the pre-metastatic niche. Cancer Cell 30, 668–681.

Maffei, M., Halaas, J., Ravussin, E., Pratley, R.E., Lee, G.H., Zhang, Y., Fei, H., Kim, S., Lallone, R., Ranganathan, S., et al. (1995). Leptin levels in human and rodent: measurement of plasma leptin and ob RNA in obese and weight-reduced subjects. Nat Med 1, 1155–1161.

Manicone, A.M., Gong, K., Johnston, L.K., and Giannandrea, M. (2016). Diet-induced obesity alters myeloid cell populations in naive and injured lung. Respir Res 17, 24.

Nagareddy, P.R., Kraakman, M., Masters, S.L., Stirzaker, R.A., Gorman, D.J., Grant, R.W., Dragoljevic, D., Hong, E.S., Abdel-Latif, A., Smyth, S.S., et al. (2014). Adipose tissue macrophages promote myelopoiesis and monocytosis in obesity. Cell Metab 19, 821–835.

Okwan-Duodu, D., Umpierrez, G.E., Brawley, O.W., and Diaz, R. (2013). Obesity-driven inflammation and cancer risk: role of myeloid derived suppressor cells and alternately activated macrophages. Am J Cancer Res 3, 21–33.

Osman, M.A., and Hennessy, B.T. (2015). Obesity correlation with metastases development and response to first-line metastatic chemotherapy in breast cancer. Clin Med Insights Oncol 9, 105–112.

Ostrand-Rosenberg, S., and Sinha, P. (2009). Myeloid-derived suppressor cells: linking inflammation and cancer. J Immunol 182, 4499–4506.

Ouzounova, M., Lee, E., Piranlioglu, R., El Andaloussi, A., Kolhe, R., Demirci, M.F., Marasco, D., Asm, I., Chadli, A., Hassan, K.A., et al. (2017). Monocytic and granulocytic myeloid derived suppressor cells differentially regulate spatiotemporal tumour plasticity during metastatic cascade. Nat Commun 8, 14979.

Paolillo, M., and Schinelli, S. (2019). Extracellular matrix alterations in metastatic processes. Int J Mol Sci 20, 4947.

Pein, M., Insua-Rodríguez, J., Hongu, T., Riedel, A., Meier, J., Wiedmann, L., Decker, K., Essers, M.A.G., Sinn, H.P., Spaich, S., et al. (2020). Metastasis-initiating cells induce and exploit a fibroblast niche to fuel malignant colonization of the lungs. Nat Commun 11, 1494.

Peinado, H., Zhang, H., Matei, I.R., Costa-Silva, B., Hoshino, A., Rodrigues, G., Psaila, B., Kaplan, R.N., Bromberg, J.F., Kang, Y., et al. (2017). Pre-metastatic niches: organ-specific homes for metastases. Nat Rev Cancer 17, 302–317.

Qian, B., Deng, Y., Im, J.H., Muschel, R.J., Zou, Y., Li, J., Lang, R.A., and Pollard, J.W. (2009). A distinct macrophage population mediates metastatic breast cancer cell extravasation, establishment and growth. PLoS One 4, e6562.

Qian, B.Z., Zhang, H., Li, J., He, T., Yeo, E.J., Soong, D.Y., Carragher, N.O., Munro, A., Chang, A., Bresnick, A.R., et al. (2015). FLT1 signaling in metastasis-associated macrophages activates an inflammatory signature that promotes breast cancer metastasis. J Exp Med 212, 1433–1448.

Quail, D.F., Olson, O.C., Bhardwaj, P., Walsh, L.A., Akkari, L., Quick, M.L., Chen, I.C., Wendel, N., Ben-Chetrit, N., Walker, J., et al. (2017). Obesity alters the lung myeloid cell landscape to enhance breast cancer metastasis through IL5 and GM-CSF. Nat Cell Biol 19, 974–987.

Rossi, E.L., de Angel, R.E., Bowers, L.W., Khatib, S.A., Smith, L.A., Van Buren, E., Bhardwaj, P., Giri, D., Estecio, M.R., Troester, M.A., et al. (2016). Obesity-associated alterations in inflammation, epigenetics, and mammary tumor growth persist in formerly obese mice. Cancer Prev Res (Phila) 9, 339–348.

Sestak, I., Distler, W., Forbes, J.F., Dowsett, M., Howell, A., and Cuzick, J. (2010). Effect of body mass index on recurrences in tamoxifen and anastrozole treated women: an exploratory analysis from the ATAC trial. J Clin Oncol 28, 3411–3415.

Singer, K., DelProposto, J., Morris, D.L., Zamarron, B., Mergian, T., Maley, N., Cho, K.W., Geletka, L., Subbaiah, P., Muir, L., et al. (2014). Diet-induced obesity promotes myelopoiesis in hematopoietic stem cells. Mol Metab 3, 664–675.

Sleeman, J.P. (2012). The metastatic niche and stromal progression. Cancer Metastasis Rev 31, 429–440.

Souma, K., Shichino, S., Hashimoto, S., Ueha, S., Tsukui, T., Nakajima, T., Suzuki, H.I., Shand, F.H.W., Inagaki, Y., Nagase, T., et al. (2018). Lung fibroblasts express a miR-19a-19b-20a sub-cluster to suppress TGF-β-associated fibroblast activation in murine pulmonary fibrosis. Sci Rep 8, 16642.

Sparano, J.A., Wang, M., Zhao, F., Stearns, V., Martino, S., Ligibel, J.A., Perez, E.A., Saphner, T., Wolff, A.C., Sledge, G.W., Jr., et al. (2012). Obesity at diagnosis is associated with inferior outcomes in hormone receptor-positive operable breast cancer. Cancer 118, 5937–5946.

Su, Y., Yao, H., Wang, H., Xu, F., Li, D., Li, D., Zhang, X., Yin, Y., and Cao, J. (2016). IL-27 enhances innate immunity of human pulmonary fibroblasts and epithelial cells through upregulation of TLR4 expression. Am J Physiol Lung Cell Mol Physiol 310, L133–141.

Sundquist, M., Brudin, L., and Tejler, G. (2017). Improved survival in metastatic breast cancer 1985-2016. Breast 31, 46–50.

Swierczak, A., and Pollard, J.W. (2020). Myeloid cells in metastasis. Cold Spring Harb Perspect Med 10, a03826.

Wang, S., Song, R., Wang, Z., Jing, Z., Wang, S., and Ma, J. (2018). S100A8/A9 in inflammation. Front Immunol 9, 1298.

Watanabe, K., Suzukawa, M., Arakawa, S., Kobayashi, K., Igarashi, S., Tashimo, H., Nagai, H., Tohma, S., Nagase, T., and Ohta, K. (2019). Leptin enhances cytokine/chemokine production by normal lung fibroblasts by binding to leptin receptor. Allergol Int 68s, S3–s8.

Williams, A., Greene, N., and Kimbro, K. (2020). Increased circulating cytokine levels in African American women with obesity and elevated HbA1c. Cytokine 128, 154989.

Wong, C.C., Gilkes, D.M., Zhang, H., Chen, J., Wei, H., Chaturvedi, P., Fraley, S.I., Wong, C.M., Khoo, U.S., Ng, I.O., et al. (2011). Hypoxia-inducible factor 1 is a master regulator of breast cancer metastatic niche formation. Proc Natl Acad Sci U S A 108, 16369–16374.

World Health Organization. (2017). Obesity and overweight fact sheet. doi:/entity/mediacentre/factsheets/fs311/en/index.html.

Yan, H.H., Pickup, M., Pang, Y., Gorska, A.E., Li, Z., Chytil, A., Geng, Y., Gray, J.W., Moses, H.L., and Yang, L. (2010). Gr-1+CD11b+ myeloid cells tip the balance of immune protection to tumor promotion in the premetastatic lung. Cancer Res 70, 6139–6149.

Yi, E.S., Bedoya, A., Lee, H., Chin, E., Saunders, W., Kim, S.J., Danielpour, D., Remick, D.G., Yin, S., and Ulich, T.R. (1996). Radiation-induced lung injury in vivo: expression of transforming growth factor-beta precedes fibrosis. Inflammation 20, 339–352.

Youn, J.I., Nagaraj, S., Collazo, M., and Gabrilovich, D.I. (2008). Subsets of myeloid-derived suppressor cells in tumor-bearing mice. J Immunol 181, 5791–5802.

Yue, X., Shan, B., and Lasky, J.A. (2010). TGF-β: Titan of lung fibrogenesis. Curr Enzym Inhib 6, 10.2174/10067.

Zaynagetdinov, R., Sherrill, T.P., Kendall, P.L., Segal, B.H., Weller, K.P., Tighe, R.M., and Blackwell, T.S. (2013). Identification of myeloid cell subsets in murine lungs using flow cytometry. Am J Respir Cell Mol Biol 49, 180–189.

