## Supplementary material for "Obesity-Activated Lung Stomal Cells Promote Myeloid-Lineage Cell Accumulation and Breast Cancer Metastasis": Figure S1, Figure S2, Table S1

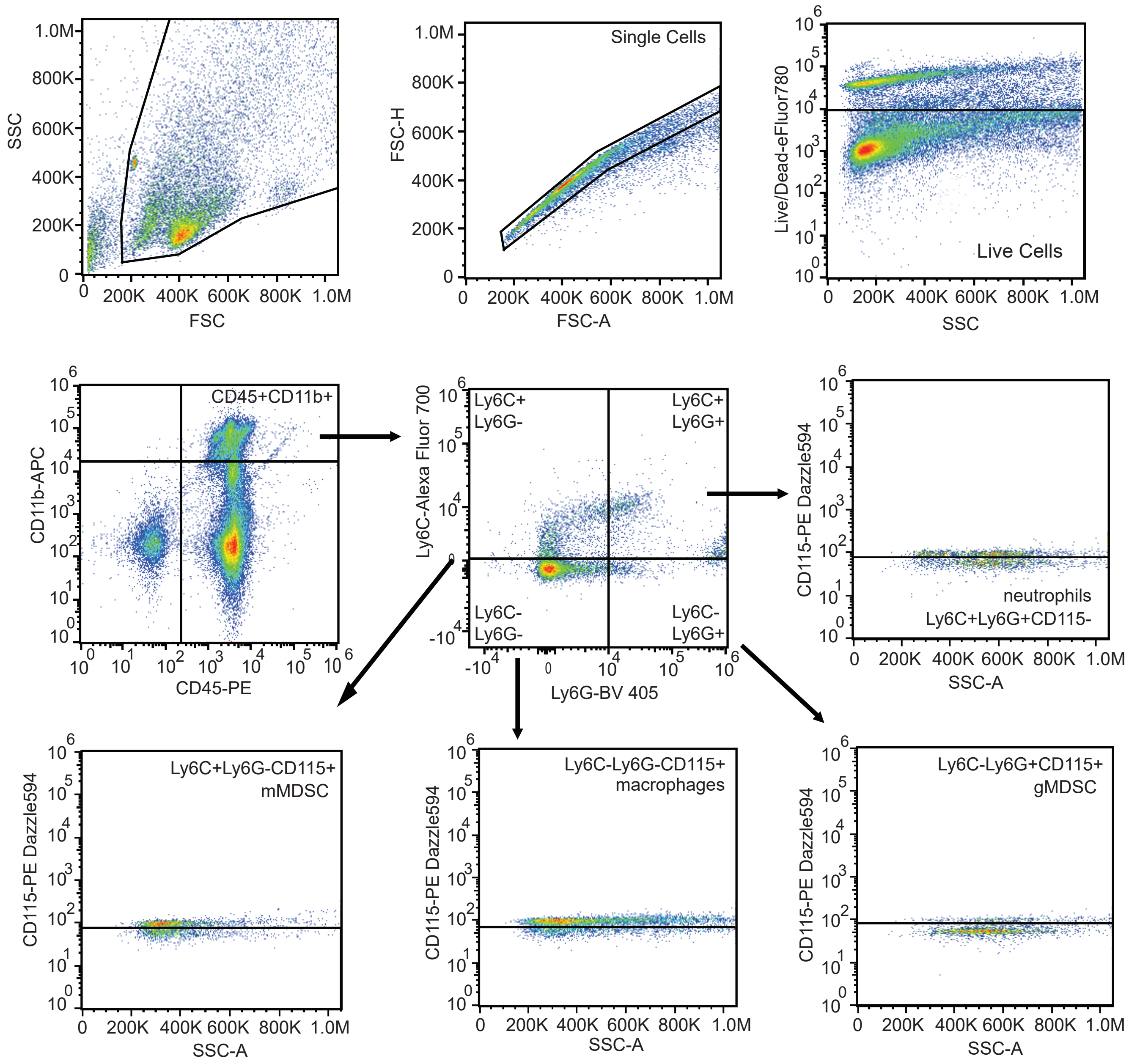


**Figure S1. Flow cytometry gating strategy of non-tumor bearing and metastatic lungs.** Debris was gated out using SSC and FCS. Doublets were removed by gating population with FSC-A and FSC-H, and live cells were identified as negative for viability dye eFluor 780. PE-conjugated CD45 and APC-conjugated CD11b were plotted to identify CD45+CD11b+ myeloid lineage cells. The CD45+CD11b+ cell population was further gated to determine Ly6C+/-Ly6G+/- populations. Each population was plotted with CD115-PE/Dazzle 594 and SSC-A to identify Ly6C+Ly6G+CD115- neutrophils, Ly6C+Ly6G-CD115+ mMDSC, Ly6C-Ly6G-CD115+ macrophages, and Ly6C-Ly6G+CD115+ gMDSC.

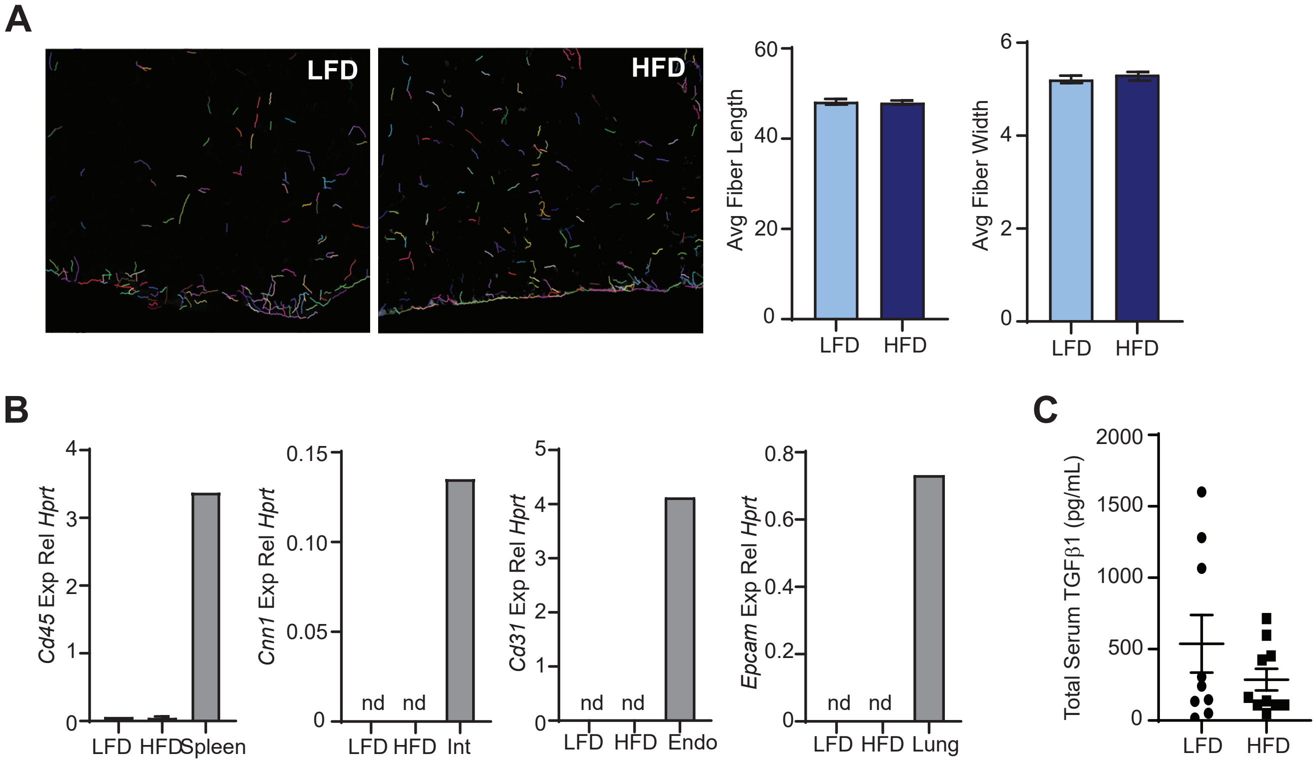


**Figure S2. Characterization of lung stromal cells from LFD or HFD-fed mice.** (A) Quantification of collagen length and width using CT-FIRE in picrosirius red-stained sections of lungs from non-tumor-bearing LFD and HFD-fed mice (n=5 mice/group). (B) Expression of markers for immune cells (*Cd45*), endothelial cells (*Cd31*), pericytes (*Cnn1* and *Cd31*), and epithelial cells (*Epcam*) from lung stromal cells isolated from LFD or HFD-fed mice. Positive controls included spleen, intestines (int), primary mouse endothelial cells (endo), and whole lung tissue. Represented as relative expression compared to *Hprt* (n=3 mice/group). (C) Concentration of total TGFβ1 protein in serum from LFD and HFD-fed mice (n=9-10 mice/group).

**Table S1. Primers used for qRT-PCR analyses.**

| | Forward 5’-3’ | Reverse 5’-3’ |
|---|---|---|
| *Hprt* | GAGTCAACGGATTTGGTCGT | GACAAGCTTCCCGTTCTCAG |
| *Csf1* | ACCCAGCTGCCCGTATGAC | TCCTTGGCAATACTCCTGCTC |
| *Csf2* | CAGGGCTGTTTTCCCATCCAT | GCCATGTTCTATCGGGTACTTC |
| *Csf3* | TGTCCTGGCCATTTCGTACC | CAGGTCTAGGCCAAGTGGTG |
| *Hif1a* | AATGCTCAGAGGAAGCGAAAAA | ATCCTTTCACTCGTTTCCAGGAA |
| *S100a8* | TGTCCTCAGTTTGTGCAGAATAAA | TTTATCACCATCGCAAGGAACTC |
| *Il1b* | GCAACTGTTCCTGAACTCAACT | ATCTTTTGGGGTCCGTCAACT |
| *Ccl2* | TTAAAAACCTGGATCGGAACCAA | GCATTAGCTTCAGATTTACGGGT |
| *Il6* | ACAAAGCCAGAGTCCTTCAGAG | GTGAGGAATGTCCACAAACTGA |
| *Il17a* | AGAAGATGCTGGTGGGTGTG | GGGTTTCTTAGGGGTCAGCC |
| *Cd31* | CTGCCAGTCCGAAAATGGAAC | CTTCATCCACCGGGGCTATC |
| *Epcam* | GAAGGGGCGATCCAGAACAA | TGTCCTTGTCGGTTCTTCGG |
| *Cd45* | GTTGTGCTTGGAGGGTCAGT | CTCAAACTTCTGGCCTTTGG |
| *Cnn1* | AACCCCACGACATCTTTGAG | AGCCAGGAGAGTGGACTGAA |
| *Col1a1* | TCTGACTGGAAGAGCGGAGA | GACGGCTGAGTAGGGAACAC |
| *Fn1* | ACGGTTTCCCATTACGCCAT | GGCACCATTTAGATGAATCGCA |
| *Lox* | CATCGGACTTCTTACCAAGCCG | GGCATCAAGCAGGTCATAGTGG |
| *Tgfβ1* | ATACGTCAGACATTCGGGAAGCAGTG | AATAGTTGGTATCCAGGGCTCTCCG |
